# Unraveling the Role of cAMP Signaling in *Giardia*: Insights into PKA-Mediated Regulation of Encystation and Subcellular Interactions

**DOI:** 10.1101/2024.04.26.591389

**Authors:** Han-Wei Shih, Germain C.M. Alas, Alexander R. Paredez

## Abstract

cAMP plays a crucial role as a second messenger in the stage transition of various protozoan parasites. This signaling pathway relies on multiple effectors, such as protein kinase A (PKA), exchange protein activated by cAMP (EPAC), and cAMP-response element binding protein (CREB) transcription factors, to initiate signal transduction in humans. Even though *Giardia* has been observed to have only two adenylate cyclases (ACs) and a single cAMP effector, the specific functions of these components are not fully understood. In our previous research, we demonstrated the essential role of AC2-dependent cAMP signaling in the upregulation of the encystation program. Using NanoBit technology, we emphasized the significance of AC2-dependent cAMP biosynthesis in regulating the dissociation of PKAr and PKAc. In this study, our objectives are twofold: first, we used newly developed Split-Halo to examine subcellular interactions of *Gl*PKAr and *Gl*PKAc; second, we investigated whether PKAc regulates encystation markers. Our findings revealed distinct subcellular locations where *Gl*PKAr and *Gl*PKAc interacted during the trophozoite stage, including the flagella, basal bodies, and cytosol. Upon exposure to encystation stimuli, the interaction shifted from the flagella to the cytosol. Additionally, interaction occurred at an unidentified cell compartment between twelve to sixteen hours of encystation and persisted to the cyst stage. Knockdown of *Gl*PKAc resulted in the downregulation of encystation-specific genes, leading to the production of fewer viable and water-resistant cysts indicating a role for PKA in transcriptional regulation. These discoveries contribute to a deeper understanding of the cAMP signaling pathway and its pivotal role in governing *Giardia*’s encystation process.

## Importance

Precise timing of interactions and subcellular compartmentation play crucial roles in signal transduction. The co-immunoprecipitation assay (CO-IP) has long been utilized to validate protein-protein interactions; however, CO-IPs lack spatial and temporal resolution. Our recent study pioneered use of the NanoBit assay, which showcased the reversible protein-protein interaction between PKAr and PKAc in response to cAMP analogs and encystation stimuli. Expanding on this groundwork, the present study employs the Split-Halo assay to uncover the subcellular compartments where PKAr and PKAc protein-protein interactions occur in response to encystation stimuli. These groundbreaking molecular tools provide a high-throughput method that combines reversibility and spatiotemporal resolution, rendering them invaluable for validating protein-protein interactions within the microbiology research field.

## Observation

*Giardia lamblia* is a protozoan parasite and the causative agent of giardiasis, a waterborne diarrheal disease that affects humans and various mammals globally. This parasite has a two-phase life cycle comprised of a resilient cyst stage and a replicative trophozoite stage. Cysts are the infectious form that transmits giardiasis. Once cysts are ingested from contaminated water and food sources, they release trophozoites that colonize the small intestine. Bile salts and an alkaline pH within the small intestine play crucial roles in stimulating the process of encystation (1). Eventually, the detached cysts transit to the lower intestine before being expelled from the host.

cAMP signaling plays a crucial role in initiating differentiation in *Giardia lamblia*, with protein kinase A (PKA) being the only identifiable conserved cAMP effector in *Giardia*. PKA activation relies on the binding of cAMP to a regulatory domain (PKAr), which liberates the catalytic kinase subunit (PKAc), rendering it active. Notably, *Giardia lamblia* has one of the smallest kinomes among eukaryotes, with only a single *Gl*PKAr (GL50803_9117) and a single *Gl*PKAc (GL50803_11214). *Gl*PKAr interacts with *Gl*PKAc in a cAMP-dependent manner (3).

Interestingly, in the trophozoite stage, *Gl*PKAr and *Gl*PKAc localize to the internal axonemes of the anterior flagella, caudal flagella, and their associated basal bodies consistent with the presence of an AKAP (3). Studies using a custom rabbit polyclonal antibody to PKA indicate that PKAr levels are diminished after the induction of encystation, while our studies of tagged PKAr show that the protein persists on internal axonemes throughout encystation (1, 3). How spatiotemporal interaction between PKAr and PKAc change during encystation are missing.

In our previous study (1), we introduced the NanoBit assay to characterize the cAMP dependent reversible interaction of *Gl*PKAr and *Gl*PKAc, providing valuable quantitative evidence of protein-protein interactions. However, while the NanoBit assay offers insights into these interactions, it lacks the ability to provide information on subcellular localization. To address this limitation, we turned to SplitHalo, an imageable reporter that has proven to be a powerful tool for investigating protein-protein interactions in human and plant cells (4, 5). In this study, we employed SplitHalo to investigate the spatiotemporal interaction between *Gl*PKAr and *Gl*PKAc in response to encystation stimuli. Additionally, we genetically knocked down *Gl*PKAc to demonstrate its significance in the encystation process.

To investigate protein-protein interactions, split fluorescent proteins have been widely utilized (7). Among these fluorescent proteins, the Halogenase (Halo) tag has proven valuable in studying protein subcellular localization due to its low background and the development of its split halo version for investigating protein-protein interactions in human and plant cells (4, 5). Because of the limited success and low fluorescence of green fluorescent protein (GFP) under anaerobic conditions, the Halo tag has also been employed as a robust fluorescent reporter for investigating protein function in anaerobic organisms (8). To precisely examine the subcellular interaction of *Gl*PKAr and *Gl*PKAc, we employed the Split-Halo assay, enabling us to observe the spatiotemporal dynamics of *Gl*PKAr and *Gl*PKAc interaction in response to encystation stimuli. The *Gl*PKAr protein (*Gl*50803_9117) was tagged with nHalo (1-155), while the *Gl*PKAc protein (GL50803_11214) was tagged with cHalo (156-257). As a control, nHalo (1-155) not fused to any protein was driven by the *Gl*PKAr native promoter (Fig 1a, SP Fig 1a).

**Fig 1.**
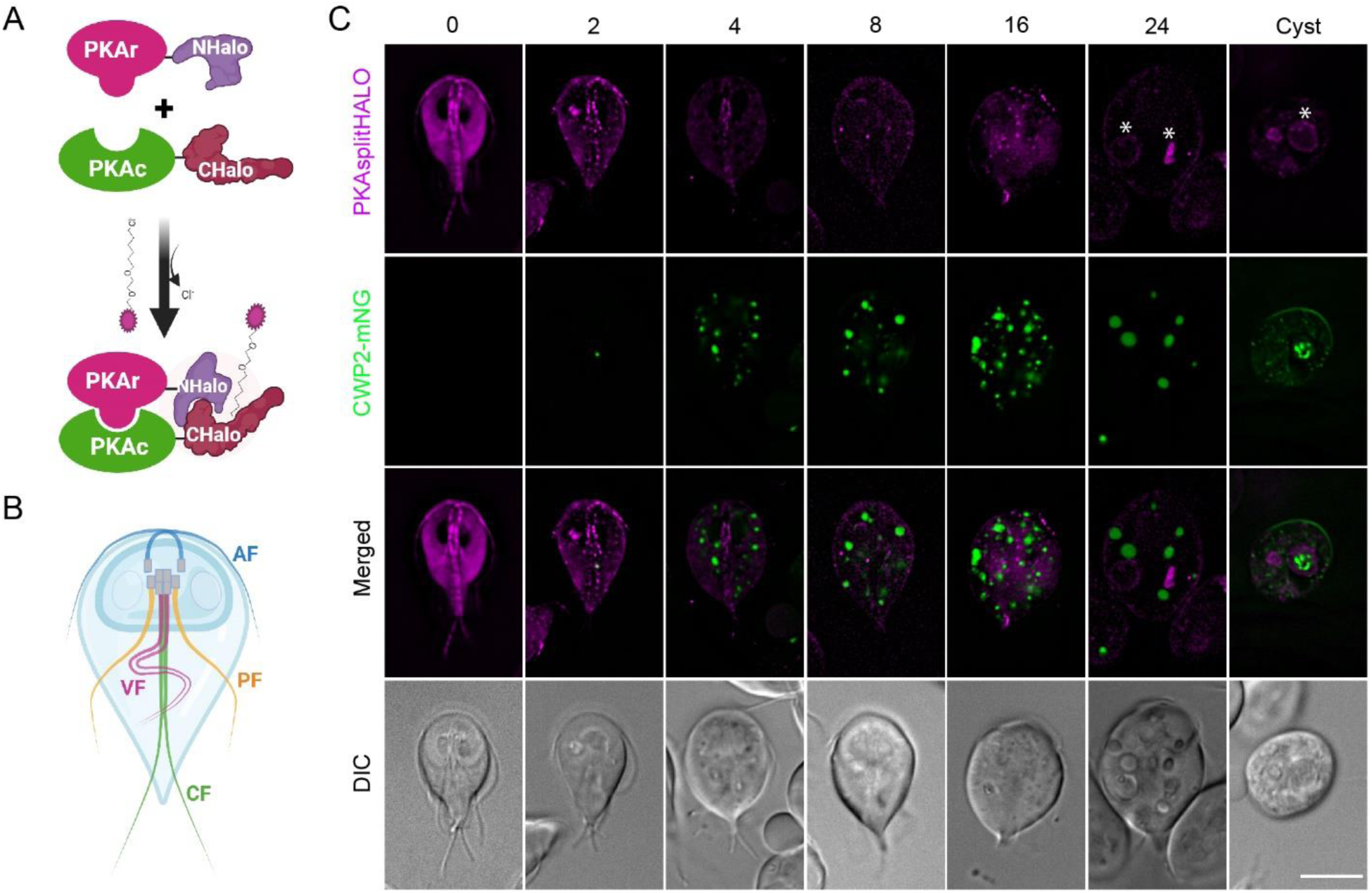
Spatiotemporal interactions of PKA-SplitHalo in response to encystation. (A) A schematic of PKA-Split Halo (GlPKAr-NHalo and GlPKAc-CHalo). Created with BioRender.com. (B) Diagram of Giardia trophozoite with labels for anterior (AF, cyan), posteriolateral (PF, yellow), caudal (CF, green), and ventral (VF, magenta) flagella. (C) Colocalization of PKA-SplitHalo with ESV marker (CWP2-mNG). Colocalization of PKA-SplitHalo and CWP2-mNG (ESV marker) at 0, 2, 4, 8, 16, 24 h and 48h (cyst) after exposure to encystation medium. White asterisk indicates the unknown compartment. Scale bars, 5 µm.

Our findings revealed specific subcellular locations where *Gl*PKAr and *Gl*PKAc interact during the trophozoite stage, including the anterior and caudal flagella, basal body, and cytosol (Fig 1b-c, SP Fig 1b). CWP2-mNeonGreen (mNG) was used as an encystation marker to observe the spatiotemporal interaction of PKA-SplitHalo in response to encystation stimuli. Upon exposure to encystation stimuli, the interaction at the anterior flagella diminished within one hour, followed by the disappearance of the interaction at the caudal flagella within two hours.

Subsequently, the primary site of interaction shifted to the cytosol from eight to twelve hours. Notably, a distinct interaction was observed at an unidentified cell compartment from twelve to sixteen hours, persisting until the cyst stage (Fig 1c).

In our prior investigation, we revealed that AC2 plays a role in the dissociation of PKAr and PKAc, leading to the upregulation of encystation-specific genes like MYB2, CWPs, and enzymes related to GalNAc biosynthesis. However, there is currently no genetic evidence supporting the involvement of *Gl*PKAc in regulating MYB2 and genes associated with cyst wall biosynthesis. To address this, we utilized CRISPRi to generate PKAc knockdown cell lines (9). The PKAc-g378, PKAc-g664, PKAc-g704, and PKAc-g848 cell lines exhibited reductions of 10%, 4%, 14%, and 35% in *Gl*PKAc-NLuc expression, respectively (SP Fig 2a).

**Fig 2.**
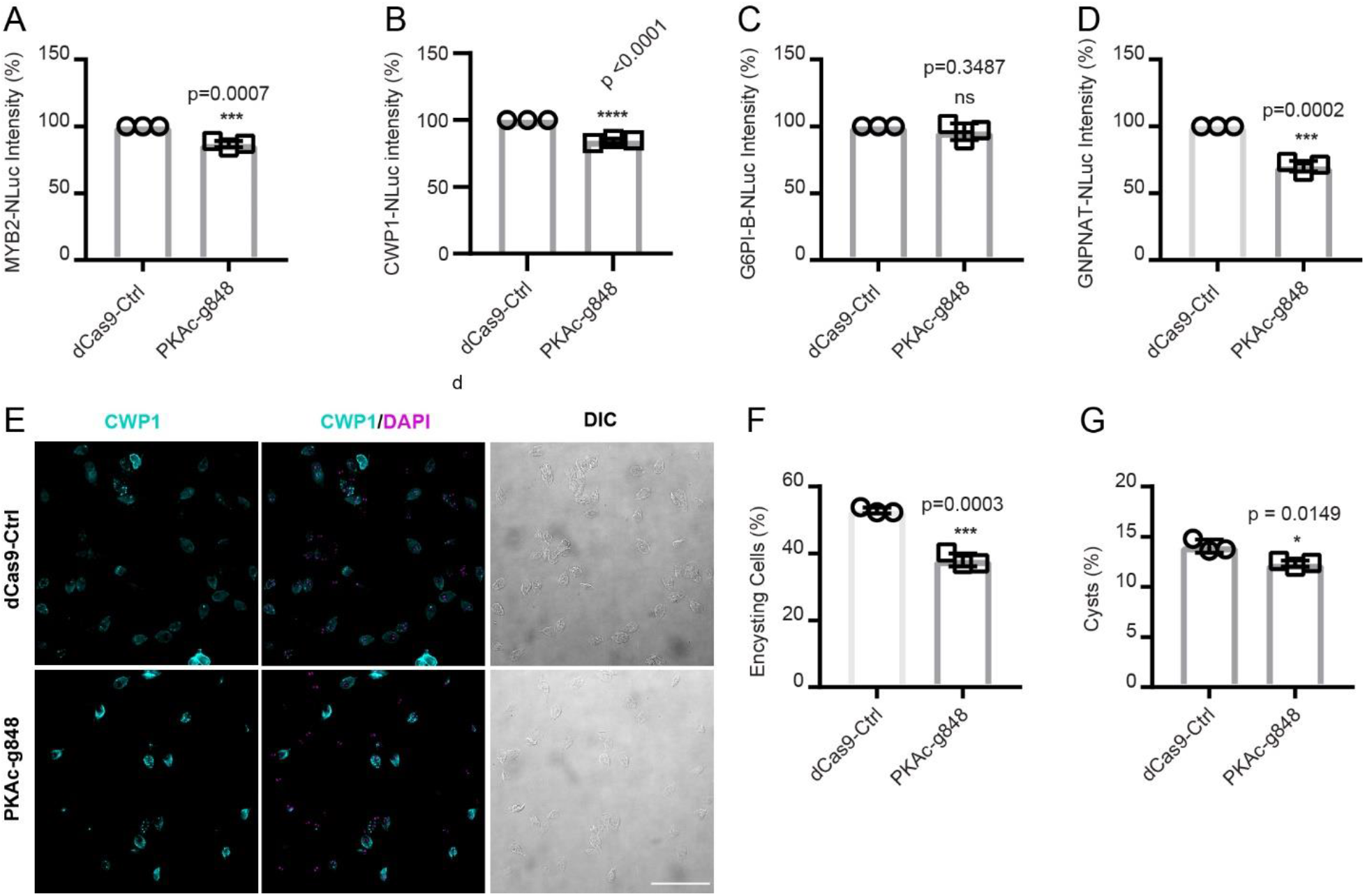
GlPKAc is critical for encystation signaling. (A-D) Relative expression levels of (A) MYB2-NLuc (GL50803_8722), (B) CWP1-NLuc, (C) G6PI-B-NLuc (GL50803_8245), (D) GNPNAT-NLuc (GL50803_14259) in dCas9-Ctrl and PKAc-g848 knockdown cell lines. (E-F) Representative IFA images (E) and quantification of 24 h encysting cells (F) from dCas9 control and PKAc-g848 knockdown cell lines (dCas9-ctrl n = 1771, and PKAc-g848 n = 1336). (G) Cyst quantification from IFA images after 24h of encystation. Scale bar, 50 µm. Data are mean ± s.d. from three independent experiments. P values were calculated with two-tailed t-tests.

We introduced PKAc-g848 into cell lines expressing encystation-specific genes with NLuc tags, including the master transcription factor (MYB2), cyst wall protein (CWP1), and two GalNAc biosynthesis enzymes (G6PI-B and GNPNAT). Upon exposure to encystation stimuli, PKAc knockdown resulted in a 15.7% decrease in MYB2 expression in the PKAc-g848 cell line (Fig 2a). Similarly, CWP1 expression decreased by 33.6% in PKAc-g848 cell lines (Fig 2b). GNPNAT-NLuc expression decreased by 30% after PKAc knockdown (Fig 2c), while G6PI-B-NLuc expression (Fig 2d) remained unaffected. Interestingly, our prior study revealed that AC2 knockdown downregulates both GNPNAT and G6PI-B expressions (1), suggesting the potential involvement of other unidentified cAMP effectors in regulating G6PI-B expression.

Furthermore, PKAc knockdown resulted in 10% and 3% reductions in encysting cells and cysts after 24 hours of exposure to encystation stimulation (Fig 2 e-g). Additionally, after 48 hours of encystation stimuli followed by 24 hours of water incubation, the PKAc knockdown cell line produced fewer water-resistant and viable cysts compared to the dCas9 control (SP Fig 2c-d). These findings underscore the crucial role of PKAc in regulating encystation signal transduction.

Additionally, we investigated the impact of the PKA inhibitor H89 on the expression of CWP1 and the number of encysting cells. Our results demonstrated that pretreatment with H89 for 1 hour reduced MYB2 and CWP1 expressions by 40 % and 33% at 4 hours post-encystation stimuli (SP Fig 3a-b) but did not affect the number of encysting cells at 24 hours (SP Fig 3 c-d). It is noteworthy that short-term treatment with H89 is limited to 1 hour, long-term exposure is lethal. We interpret this result to indicate that by 24 hours the treated cells are able to catch up to untreated control cells.

In summary, we employed the Split-Halo assay to investigate subcellular interactions between *Gl*PKAr and *Gl*PKAc in *Giardia*. Spatiotemporal changes in the interaction were observed in response to encystation stimuli, with shifts in cellular locations over time. Further exploration of the unidentified cell compartment and the dynamics of *Gl*PKAr and *Gl*PKAc interaction during encystation could reveal novel mechanisms involved in cyst formation. CRISPRi-mediated knockdown of *Gl*PKAc resulted in altered expression of encystation-specific genes, including MYB2, CWP1, and GNPNAT, highlighting *Gl*PKAc’s role in encystation signal transduction and regulation of transcriptional responses to encystation stimuli. This genetic manipulation also led to reduced encysting cell numbers and impaired cyst viability. These findings confirm the utility of Split-Halo for protein-protein interactions and the role of PKA in regulating encystation.

Future efforts will focus on identifying the phosphorylation targets of PKA.

## Material and methods

### Design of guide RNA for CRISPRi

Guide RNA for the CRISPRi system utilized the Dawson Lab protocol (9), NGG PAM sequence and *G. Lamblia* ATCC 50803 genome were selected for CRISPRi guide RNA design with Benchling. PKAc-NLuc was created from previous study. Primers are listed in table 1.

### Design of GlPKA-SplitHalo

The PKA-NanoBit system and its control, as established in a previous study, were employed to create GlPKA-SplitHalo. In summary, the cHalo fragment (156-257) was PCR amplified from Halo7 and ligated with PKAc (GL50803_101214) under its native promoter (∼500 bp), while the nHalo fragment (1-155) was amplified and attached to PKAr (GL50803_9117) also under its native promoter (∼500 bp) (Fig. 1a and Supplementary Fig. 1a). The control cell line expresses PKAc-cHalo (156-257) and the nHalo fragment (1-155) under the native promoter of PKAr (pPKAr), without being fused to any protein (see Supplementary Fig. 1a). Primers are listed in table 1.

### *Giardia* growth and encystation media

The *Giardia intestinalis* isolate WB clone C6 (American Type Culture Collection catalog number 50803;) was cultured in Keister’s modified TYI-S33 media supplemented with 10% adult bovine serum and 0.125 mg/ml bovine bile at pH 7.1. To induce encystation, cells were initially incubated for 48 hours in pre-encystation media without bovine bile at pH 6.8, followed by further incubation with media at pH 7.8 supplemented with 10% adult bovine serum, 0.25 mg/ml porcine bile (B8631, Sigma-Aldrich), and 5 mM calcium lactate. H89 inhibitor treatments, specifically N-[2-(p-Bromocinnamylamino) ethyl]-5-isoquinolinesulfonamine dihydrochloride (H89, B1427, Sigma-Aldrich), were utilized in this study.

### Live imaging

For live imaging, parasites were released from culture tubes by chilling and then were transferred to Attoflor imaging chambers that were incubated for 60 min with or without treatments in a Panasonic trigas incubator set to 5% CO2 and 2% O2. To detect HALO tag, 20 nM Janelia Fluor® HaloTag® Ligand (Promega, GA112A) was used. Before imaging, parasites were washed with 1XHBS to remove detached cells and excess Halo ligand. Images were acquired on a DeltaVision Elite microscope using a 60X, 1.42-numerical aperture objective with a PCO Edge sCMOS camera, FITC and Cy5 filter set, and images were deconvolved using SoftWorx (API, Issaquah, WA).

### *In vitro* bioluminenscence assays

*Giardia* cells were chilled on ice for 15 minutes and then centrifuged at 700 x g for 7 minutes at 4°C. Subsequently, the cells were resuspended in cold 1X HBS (HEPES-buffered saline), and dilutions were prepared after determining cell density using a MOXI Z mini Automated Cell Counter Kit (Orflo, Kenchum, ID). For NanoLuc luminescence measurement, 20,000 cells were loaded into white polystyrene, flat-bottom 96-well plates (Corning Incorporated, Kennebunk, ME) and mixed with 10 μl of NanoGlo luciferase assay reagent (Promega). Relative luminescence units (RLU) were detected using a pre-warmed 37°C Perkin Elmer Victor-3 1420 Multilabel Counter for 30 minutes to achieve the maximum value. These experiments were conducted with three independent bioreplicates.

### Immunofluorescence

Giardia parasites were chilled on ice for 30 minutes and then centrifuged at 700 x g for 7 minutes. The resulting pellet was fixed in PME buffer (100 mM Piperazine-N,N’-bis(ethanesulfonic acid) (PIPES) pH 7.0, 5 mM EGTA, 10 mM MgSO4) supplemented with 1% paraformaldehyde (PFA) (Electron Microscopy Sciences, Hatfield, PA), 100 µM 3-maleimidobenzoic acid N-hydroxysuccinimide ester (Sigma-Aldrich), 100 µM ethylene glycol bis(succinimidyl succinate) (Pierce), and 0.025% Triton X-100 for 30 minutes at 37°C. The fixed cells were then attached to polylysine-coated coverslips. After washing once with PME, the cells were permeabilized with 0.1% Triton X-100 in PME for 10 minutes. Following two quick washes with PME, blocking was carried out in PME supplemented with 1% bovine serum albumin, 0.1% NaN3, 100 mM lysine, and 0.5% cold water fish skin gelatin (Sigma-Aldrich). Subsequently, the cells were incubated with a 1:200 dilution of Alexa 647-conjugated anti-CWP1 antibody (Waterborne, New Orleans, LA) for 1 hour. After three washes in PME plus 0.05% Triton X-100, coverslips were mounted with ProLong Gold antifade plus 4’,6-diamidino-2-phenylindole (DAPI; Molecular Probes). Images were captured using a DeltaVision Elite microscope equipped with a 60X, 1.42 numerical aperture objective and a PCO Edge sCMOS camera, followed by deconvolution using SoftWorx (API, Issaquah, WA).

### Image analysis

We employed ImageJ for processing all images, and figures were assembled using Adobe Illustrator. Cyst count and cyst viability staining *Giardia* trophozoites were cultured for 24 hours in encystation media supplemented with 10 g/L ovine bovine bile and calcium lactate, followed by another 24 hours in TYI-S33 media. After a total of 48 hours, the total cell number was determined using a MoxiZ Coulter counter. The encysted culture was then centrifuged at 700 x g for 7 minutes, and the pellets were washed 10 times with deionized water before being stored in distilled water overnight at 4°C. To determine cyst concentration, 20 μl of the 48-hour encysted cells were counted using a hemocytometer. Cyst viability was assessed using fluorescein diacetate (FDA) and propidium iodide (PI) staining to differentiate live and dead cysts. Images were captured using a DeltaVision Elite microscope equipped with a 40X, 1.35 numerical aperture objective and a PCO Edge sCMOS camera. The images were subsequently deconvolved using SoftWorx (API, Issaquah, WA).

---

**SP Fig 1.**
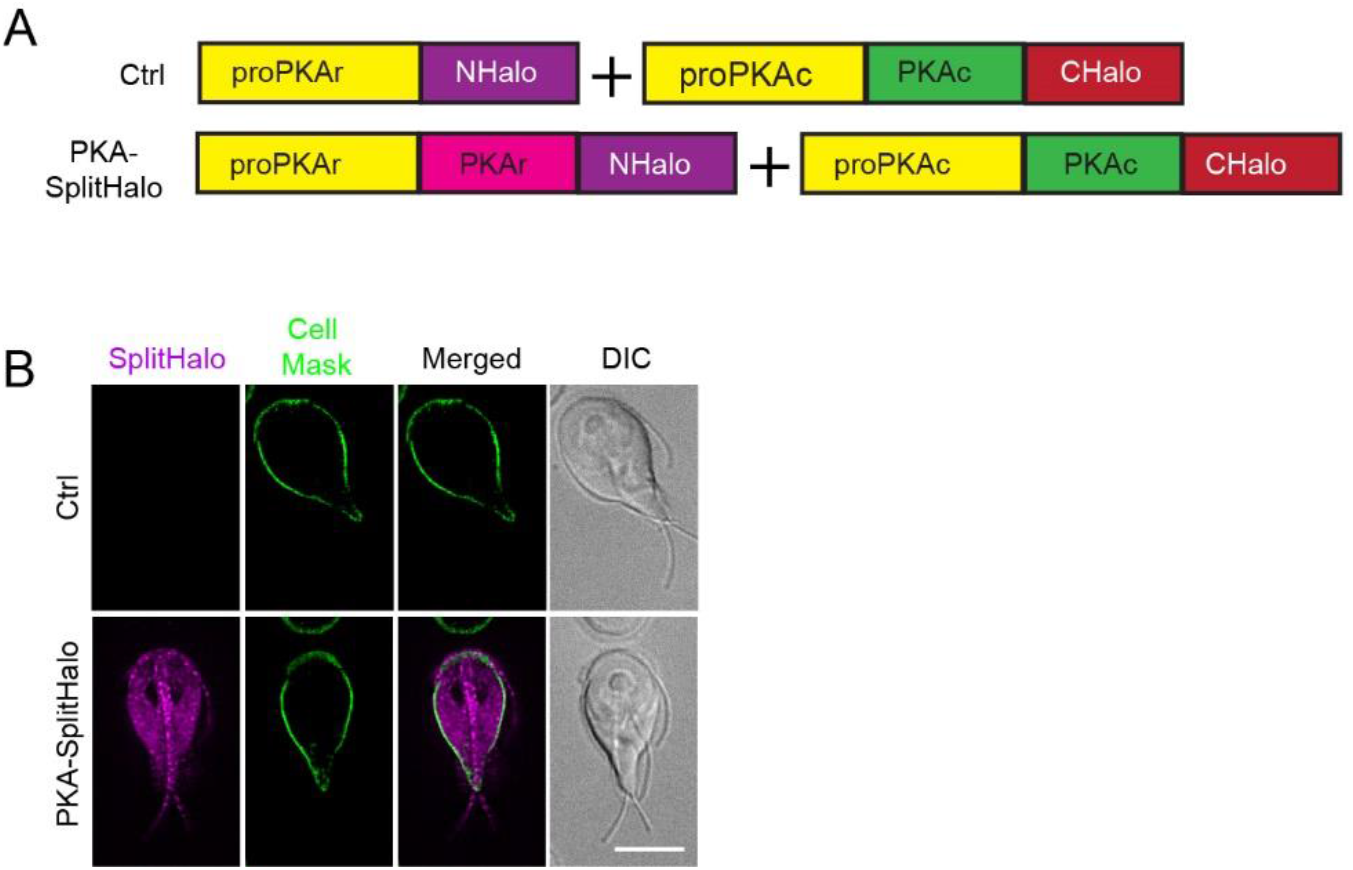
Design of GlPKA-SplitHalo. (A) GlPKA-Split Halo is composed of pPKAr::PKAr-nHalo (1-155) and pPKAc::PKAc-cHalo (156-257), and the control is composed of pPKAr::nHalo and pPKAc::PKAc-cHalo. (B) Localizations of PKA-SplitHalo and control cell lines. Cell mask is used to mark plasma membrane. Scale bars, 5 µm.

**SP Fig 2.**
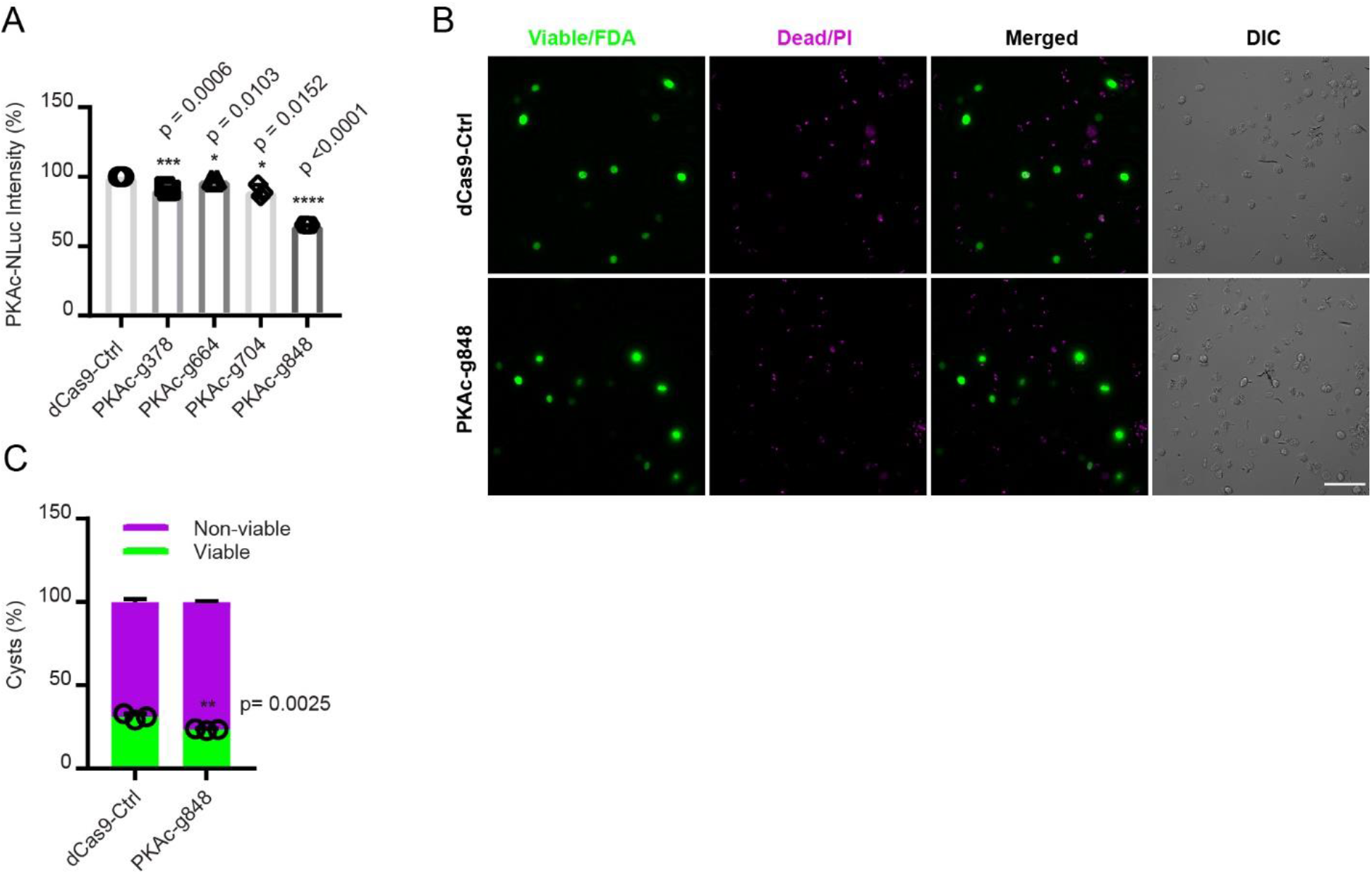
Cyst viability of PKAc knockdown cell lines. (A) Screening of PKAc guide RNAs. Relative PKAc-NLuc levels using the indicated CRISPRi gRNAs. (B-C). Representative images (B) and quantification (C) of dCas9 control and PKAc-g848 derived cysts stained with fluorescein diacetate (FDA, green=live) and propidium iodine (PI, magenta=dead). dCas9-ctrl n=752, and PKAc-g848 n=641 Scale bar, 50 µm. Data are mean ± s.d. from three independent experiments. P values were calculated with two-tailed t-tests.

**SP Fig 3.**
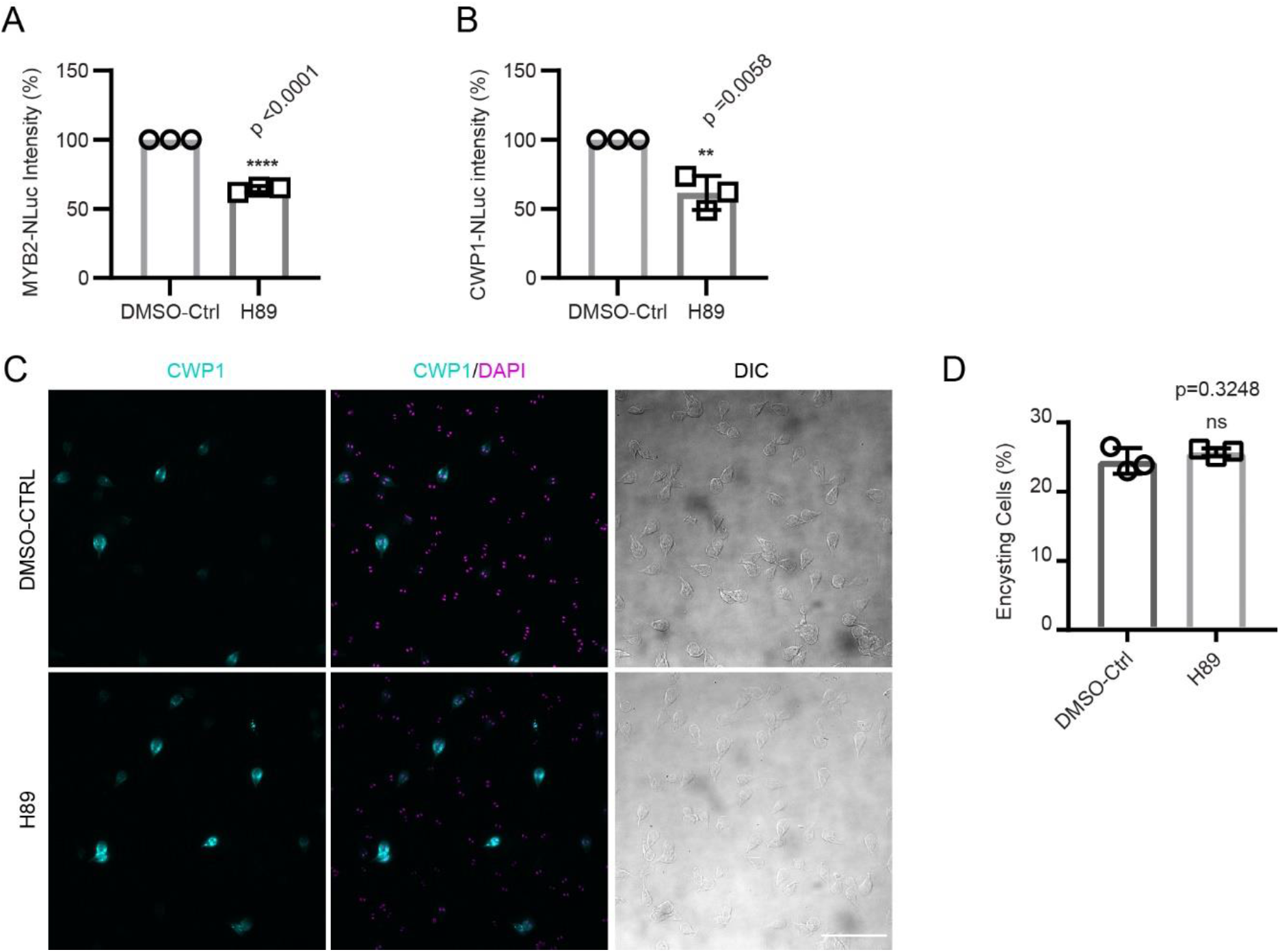
PKAc inhibitor H89 impairs encystation. (A-B) Relative expression levels of (A) MYB2-NLuc (GL50803_8722) and (B) CWP1-NLuc in DMSO or H89 (1h pretreatment). (C-D) Representative IFA images (C) and quantification of 24 h encysting cells (D) from 1h pretreatment with DMSO or H89. Parasites were exposed to encystation medium for 24 h and stained with CWP1 antibody and DAPI. Scale bar, 50 µm. Data are mean ± s.d. from three independent experiments. P values were calculated with two-tailed t-tests. (total cells counted for DMSO-Ctrl n= 1002, and H89 n=1028).

